# HCV- and HBV-mediated liver cancer converge on similar transcriptomic landscapes and immune profiles

**DOI:** 10.1101/2024.07.01.601493

**Authors:** Elizabeth S. Borden, Annika Jorgensen, Heini M. Natri, Karen Taraszka Hastings, Kenneth H. Buetow, Melissa A. Wilson

## Abstract

Hepatocellular carcinoma (HCC) remains a leading cause of cancer-related deaths worldwide, and a large proportion of HCC is attributable to viral causes including hepatitis B (HBV) and C virus (HCV). The pathogenesis of viral-mediated HCC can differ between HBV and HCV, but it is unclear how much these differences influence the tumors’ final molecular and immune profiles. Additionally, there are known sex differences in the molecular etiology of HCC, but sex differences have not been explored in the context of viral-mediated HCC. To determine the extent to which the viral status and sex impact the molecular and immune profiles of HCC, we performed differential expression and immune cell deconvolution analyses. We identified a large number of differentially expressed genes unique to the HBV or HCV tumor:tumor-adjacent comparison. Pathway enrichment analyses demonstrated that the changes unique to the HCV tumor:tumor-adjacent tissue were predominated by changes in the immune pathways. Immune cell deconvolution demonstrated that HCV tumor-adjacent tissue had the largest immune cell infiltrate, with no difference in the immune profiles within HBV and HCV tumor samples. We subsequently segregated the differential expression analyses by sex, but demonstrated that the low number of female samples led to an overestimate of differentially expressed genes unique to male tumors. This limitation highlights the importance of additional sampling of female HCC tumors to allow for a more complete analysis of the sex differences in HCC. Overall, this work demonstrates the convergence of HBV- and HCV-mediated HCC on a similar transcriptomic landscape and immune profile despite differences in the surrounding tissue.

**Author Summary:** Hepatocellular carcinoma (HCC) is a significant worldwide health challenge. The majority of cases are attributable to infection with hepatitis B (HBV) or C (HCV). HBV and HCV differ in their methods of transmission and how they cause cancer. Despite these differences, most treatment guidelines are ambivalent to the underlying viral etiology of the tumor. In the age of personalized medicine, we sought to determine how similar or different HCC was when mediated by HBV or HCV. Additionally, since previous work has demonstrated biological differences in the tumors between males and females, we sought to characterize the sex differences within viral-mediated HCC. We found that, although there are several genes with differences in HBV- and HCV-mediated tumors, the tumors appear to be more biologically similar than the corresponding tumor-adjacent tissue. This suggests a convergence on common pathways toward cancers even when the starting point differs. The lower number of female samples inhibits a full understanding of the biological differences between HCC in males and females. This presents a critical need in the field to increase the sampling of female cancers to enable a full understanding of the sex differences in HCC.

## Introduction

Hepatocellular carcinoma (HCC) remains a critical health challenge worldwide, leading to over 600,000 deaths annually (1). Risk factors for HCC include hepatitis B, C, and D viruses (HBV, HCV, and HDV); alcoholic liver disease; and non-alcoholic fatty liver disease (2). Approximately 50% of HCC is attributable to HBV infection (2). The risk of HCC with HCV has been reduced with the introduction of antiviral therapies that have led to a sustained virological response to HCV (3). However, antiviral therapies do not reduce the risk of HCC in patients who have already progressed to cirrhosis, and 30% of HCC remains attributable to HCV infection (1).

The pathogenesis of HCC from HBV and HCV can be attributable to multiple underlying mechanisms. Both viruses can be associated with persistent inflammation, immune-mediated oxidative stress from chronic infection, and abnormal regulation of signaling pathways (1,4). HBV can integrate into the host genome causing insertional mutagenesis, which may be carcinogenic (5). Chronic HCV has also been associated with steatosis and subsequent progression to fibrosis and cirrhosis (6). Additionally, in endemic areas, HBV is primarily transmitted perinatally (7), leading to lifelong infections that may elicit significantly different immune responses compared to HCV which is primarily transmitted through direct contact with blood later in life (8).

Despite the differences in the underlying pathogenesis of viral-mediated HCC, the majority of treatment guidelines do not discriminate based on etiology (9–11). Within the last few years, immune checkpoint inhibitors, including anti-PD1, anti-PDL1, and anti-CTLA-4 treatments, have been FDA-approved for the treatment of advanced HCC (12–14). Studies to date have been underpowered to detect differences in the response of HBV and HCV to immune checkpoint inhibitors (15,16). Previous studies have demonstrated differences in the immune microenvironment in HBV- and HCV-mediated HCC (17), suggesting the importance of understanding the tumor microenvironment in viral-mediated liver cancer.

Sex differences in the incidence, mortality, and genetic profile of HCC have been documented (18,19), and previous work from our lab has documented sex differences in the molecular etiologies of HCC (20). However, there is less known about the sex differences in viral-mediated HCC. Here, we performed differential expression analyses on a cohort of viral-mediated HCC cases with paired tumor-adjacent tissue. We segregated our analyses first by viral etiology and then by the combination of viral etiology and sex to illuminate the underlying molecular profiles and immune landscapes in these tumors and the adjacent tissue. We discuss the challenges stemming from sampling biases and the need for increased sampling of female tumors to fully probe the biological mechanisms leading to the differences in incidence and mortality from HCC in males and females. Furthermore, while we identified several genes with differential expression in HBV- and HCV-mediated liver cancer, we demonstrated that the tumor tissue appears to converge on a more similar transcriptional landscape and immune profile compared to the tumor-adjacent tissue. Together these results highlight the importance of considering sex and etiology in defining the transcriptional and immune profiles of HCC.

## Results

### Segregation by viral etiology enables the identification of distinct sets of differentially expressed genes

HBV and HCV are both known etiologies for HCC, despite their differences in viral class, genome type, and transmission route (Figure 1A). To probe the molecular phenotypes of HCC arising from HBV or HCV, we performed differential expression analyses across all tumor vs. tumor-adjacent samples regardless of etiology (147 tumor vs. 129 tumor-adjacent, Figure 1B), HCV tumor vs. tumor-adjacent samples only (102 tumor vs. 91 tumor-adjacent, Figure 1C), and HBV tumor vs. tumor-adjacent samples only (45 tumor vs. 38 tumor-adjacent, Figure 1D). Genes were considered differentially expressed if they had a false-discovery rate of less than 0.05.

**Figure 1:**
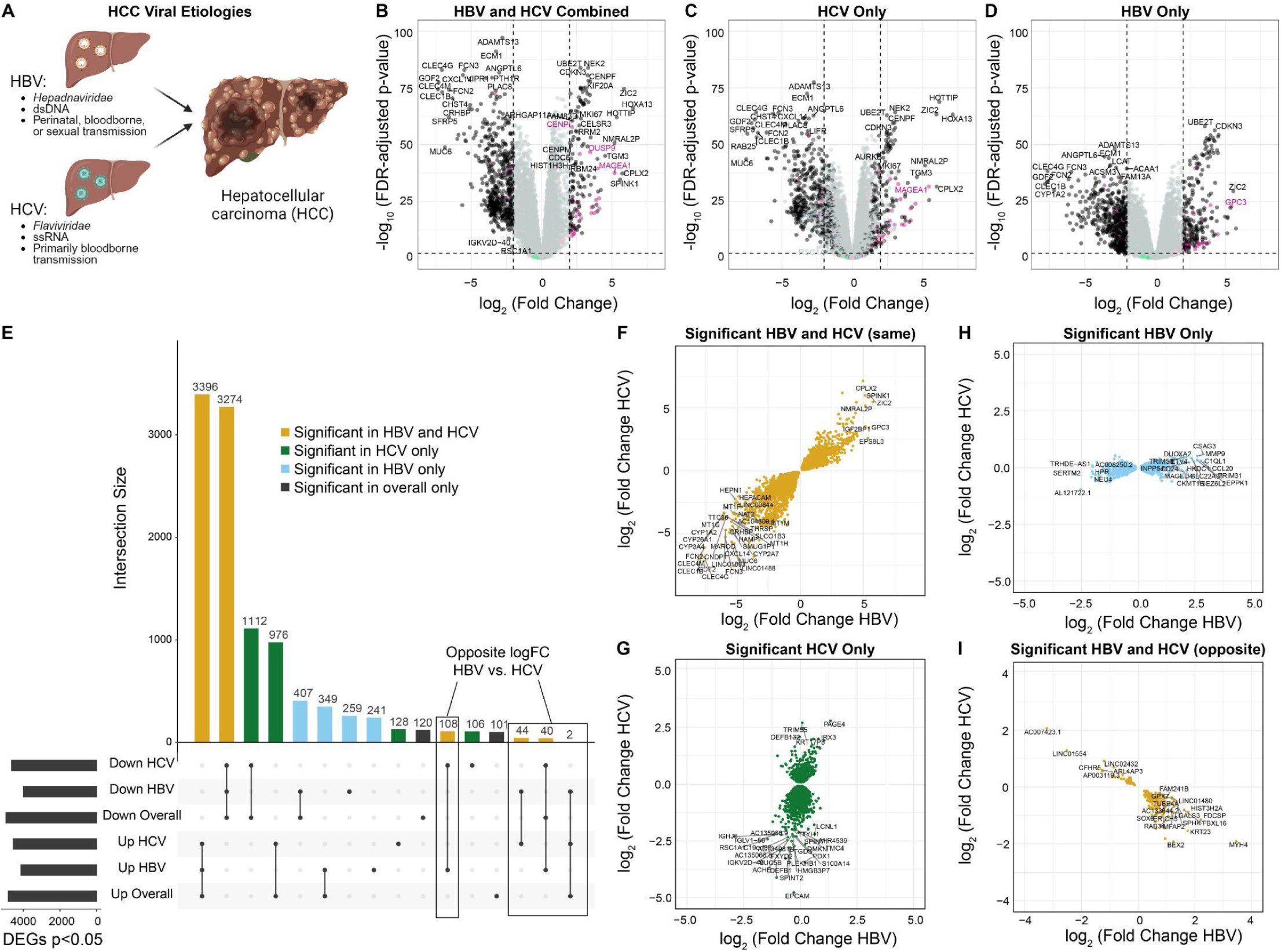
Segregation by viral etiology increases the identification of differentially expressed genes. **(A)** Overview of hepatitis B (HBV) and hepatitis C (HCV). HBV is a *hepadnaviridae* with a double-stranded DNA (dsDNA) genome and is primarily transmitted perinatally, but can also be bloodborne or sexually transmitted. HCV is a *flaviviridae* with a single-stranded RNA (ssRNA) genome and has primarily bloodborne transmission. **B-D)** Volcano plots of differentially expressed genes from **(B)** HBV and HCV combined, **(C)** HCV only, and **(D)** HBV only. X-linked genes are indicated in pink, Y-linked in green, and autosomal in black. **E)** Upset plot of differentially expressed genes from each of the comparisons. Genes shared in all comparisons are colored yellow, genes unique to the HCV subset are green, and genes unique to the HBV subset are light blue. **F-I)** Comparison of the logFC in HBV and HCV for **(F)** genes significant in both HBV and HCV tumors in the same direction, **(G)** genes significant only in the HCV subset, **(H)** genes significant only in the HBV subset, and **(I)** genes significant in both HBV and HCV but in opposite directions.

We then evaluated the degree of overlap between the lists of differentially expressed genes from each comparison (Figure 1E). Across all comparisons, we detect 10,663 unique differentially expressed genes. Of these, 6,670 (62.6%, 3,396 upregulated in tumors and 3,274 downregulated in tumors) were significant across all samples, HCV samples, and HBV samples. 2,088 genes (19.6%, 1,112 downregulated in tumors, 978 upregulated in tumors) were identified as significant in the comparison across all samples as well as HCV samples alone, but not in the analysis of HBV samples alone. Similarly, 756 genes (7.1%, 407 downregulated in tumors, 349 upregulated in tumors) were significant in the analysis of all samples and HBV samples alone, but not in HCV samples alone. Moreover, 500 genes not identified as differentially expressed in the overall comparison were identified as differentially expressed in HBV samples only (259 down-regulated in tumors and 241 up-regulated), while 234 genes not identified as differentially expressed in the overall comparison were identified as differentially expressed in HCV samples (128 up-regulated in tumors and 106 down-regulated). 221 genes identified in the overall comparison were not detected as significant in either HBV or HCV samples when segregated.

Finally, 194 genes were identified as differentially expressed in opposite directions in HBV and HCV samples. Of these, 152 were not identified in the overall comparison but were differentially expressed in opposite directions in HBV and HCV when segregated (108 down-regulated in HCV tumor samples and up-regulated in HBV tumor samples, and 44 down-regulated in HBV tumor samples and up-regulated in HCV tumor samples). 42 genes were identified as differentially expressed in the overall comparison and then identified in opposite directions in HBV and HCV when segregated. Of the 42, 40 were originally identified as down-regulated in all tumor samples and then identified as down-regulated in HCV tumor samples and up in HBV tumor samples when segregated, and 2 were originally identified as up-regulated across all tumor samples and then up in HCV tumor samples and down in HBV tumor samples when segregated.

We then evaluated whether the genes identified in the segregated analyses showed similar expression changes from tumor-adjacent to tumor tissue in HBV and HCV samples, despite not passing multiple testing corrections in both etiologies. We plotted the log fold change (logFC) of the tumor:tumor-adjacent change in HBV compared to HCV for each subset of genes (Figure 1F-I). Across the genes with shared differential expression in HBV and HCV, we fit a linear model and found that our model had an R-squared value of 0.833, suggesting that 83.3% of the variability in the data is explained through the model (Figure 1F). By contrast, for the differentially expressed genes expressed in HCV only, the R-squared value is 0.200 (Figure 1G) and for the genes differentially expressed in HBV only, the R-squared value is only 0.197 (Figure 1G). This finding supports that the differences observed are more likely to be biological differences in the transcriptional profiles of HBV- and HCV-mediated HCC rather than artifacts of the differences in sample size. A majority (95.4%) of the differentially expressed genes with opposite directions in HBV and HCV had an absolute logFC of less than 1.5 in both HBV and HCV. However, a small number show large opposite-fold changes, including *BEX2* (24,25), *AP1M2* (26), and *KRT23* (27,28), all of which have been previously associated with the pathogenesis of HCC (Figure 1I).

### Genes unique to HCV are enriched for immune pathways

To identify pathways enriched in the differentially expressed genes, we performed gene ontology enrichment analyses on the shared and unique differentially expressed genes for HBV and HCV (Figure 2). The pathways enriched in the shared differentially expressed genes are consistent with the hallmarks of cancer. These pathways include the immune response (“Immunoglobulin Mediated Immune response”), cell division (“Mitotic Nuclear division” and “Regulation of Mitotic Nuclear Division”), and the “Epoxygenase P450 Pathway” which is known to regulate the hepatic inflammatory response (Figure 2A). Within the genes uniquely differentially expressed in HCV, we identified a predominance of immune-related pathways, including “Regulation of T cell Activation”, “Response to Virus”, “Antigen processing and presentation”, “MHC protein complex assembly”, and “Leukocyte cell adhesion” (Figure 2B). Finally, in the genes significant to HBV only, a few pathways were identified, predominantly the “Intrinsic apoptotic signaling pathways” (Figure 2C).

**Figure 2:**
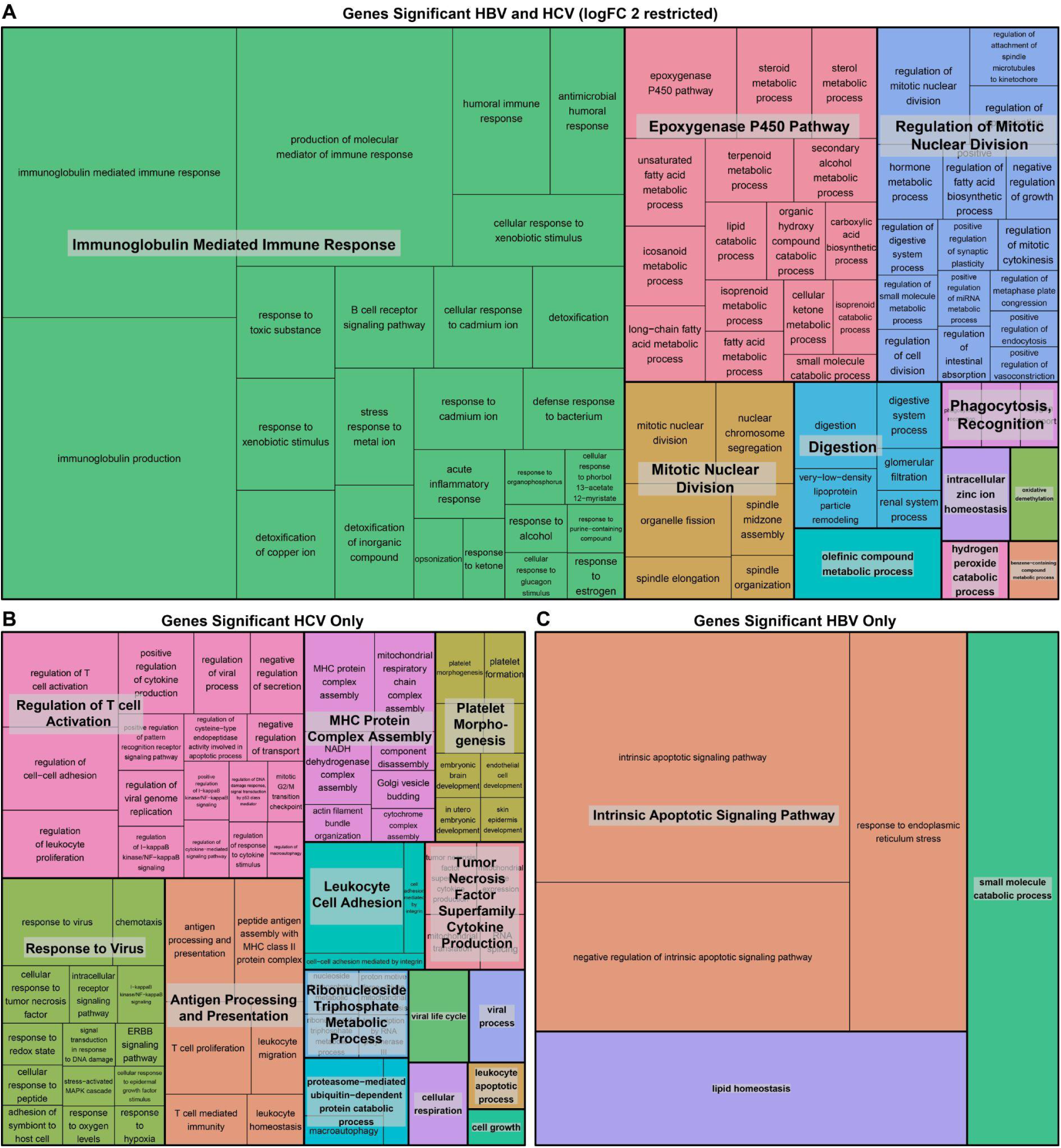
Genes unique to HCV tumor:tumor-adjacent comparison are enriched for immune pathways. Treemap visualization of GO enrichment analysis from **(A)** overall tumor vs. tumor-adjacent comparison (restricted by a logFC > 2 due to the large number of significant genes), **(B)** genes unique to HCV tumor vs tumor-adjacent comparison, and **(C)** genes unique to HBV tumor vs. tumor-adjacent comparison. The sizes of the boxes reflect the magnitude of the false-discovery adjusted p-value for the GO enrichment term.

### Tumor-adjacent tissue is more distinct than tumor tissue based on viral etiology

While the previous analyses identified genes that change expression from tumor-adjacent to tumor tissue, stratified by etiology, we were interested in directly examining etiology-based differences in tumor-adjacent and tumor tissue. We therefore performed differential expression analyses on HBV tumor vs. HCV tumor and HBV tumor-adjacent vs. HCV tumor-adjacent samples. When differentially expressed genes are defined as genes with an FDR < 0.05, 2,363 genes were identified as upregulated in HCV tumor-adjacent tissue and 1,817 upregulated in HBV tumor-adjacent tissue. By contrast, a smaller number of genes are found to be differentially expressed in the tumor tissue with 341 upregulated in HCV and 364 upregulated in HBV (Figure 3A-B). Of note, when restricting to the genes with the largest logFCs (>2), 14/21 genes upregulated in HCV tumor-adjacent tissue were immunoglobulin genes and 2/21 were interferon-inducible genes, indicating a strong predominance of immune-related genes (Supplementary Figure 1A-U). To establish whether these genes are normally expressed in healthy liver tissue, we probed the expression of each across the normal liver GTEx samples. The majority of genes upregulated in HCV tumor-adjacent tissue, except *IFI27*, *IFI6*, and *PLA2G2A*, show average expression below 1 TPM in GTEx normal liver samples, suggesting that the upregulation of these immune-related genes in HCV tumor-adjacent tissue is likely mediated by the viral infection (Supplementary Figure 1V).

**Figure 3:**
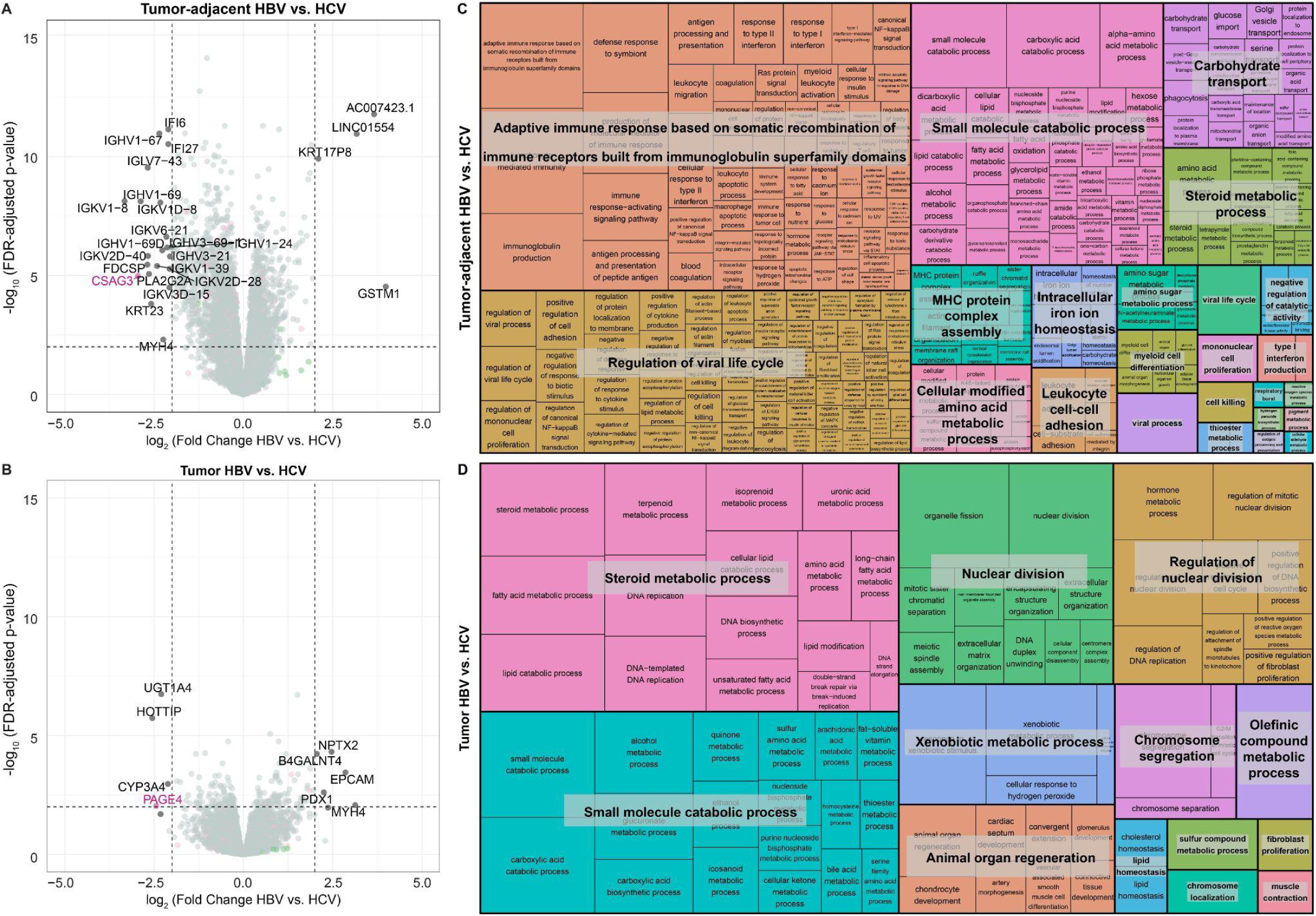
Tumor-adjacent tissue is more distinct than tumor tissue based on viral etiology. Volcano plots of differentially expressed genes from **(A)** tumor-adjacent HBV vs. HCV and **(B)** tumor HBV vs. HCV samples. In each comparison, genes with higher expression in HCV are on the left side of the plot with negative log fold changes and genes with higher expression in HBV are on the right side of the plot with positive fold changes. Genes with a false discovery rate (FDR) < 0.05 and a log fold change (logFC) > 2 are labeled on the plot. X-linked genes are indicated in pink, Y-linked in green, and autosomal in black. Treemap visualization of GO enrichment analysis from **(A)** tumor-adjacent HBV vs. HCV and **(B)** tumor HBV vs. HCV. The sizes of the boxes reflect the magnitude of the false-discovery adjusted p-value for the GO enrichment term.

We then performed pathway enrichment analysis on all differentially expressed genes between HBV and HCV in both the tumor and tumor-adjacent samples. Within the HBV tumor-adjacent:HCV tumor-adjacent comparison, pathway enrichment analysis demonstrated enrichment of immune-related pathways including “Adaptive immune response based on somatic recombination of immune receptors built from immunoglobulin superfamily domains”, “Regulation of viral life cycle”, and “MHC protein complex assembly” (Figure 3C). By contrast, within the smaller number of differentially expressed genes in the tumor tissue, the primary differences were found in metabolism (“Steroid metabolic process”, “Small molecule catabolic process”, and “Xenobiotic metabolic process”) and nuclear division (“Nuclear division”, “Regulation of nuclear division”, and “Chromosome segregation”) (Figure 3D).

Given the predominance of immune-related pathways identified in the 1) HCV tumor:HCV tumor-adjacent and 2) HBV tumor-adjacent:HCV tumor-adjacent comparisons, we performed immune cell deconvolution across all samples. Immune cell deconvolution was first performed with xCell (29) and demonstrated a significant increase in the immune cell infiltration in HCV tumor-adjacent samples compared to all other samples. This increase is seen in the overall immune score, accounting for all immune cell populations (Figure 4A, Supplementary Figure 2A). Since the enriched pathways suggested changes in the adaptive immune response specifically, we examined the CD8+ T cell population and demonstrated the highest infiltration of CD8+ T cells in HCV tumor-adjacent samples (Figure 4B, Supplementary Figure 2A). Despite the increased immune infiltration in HBV and HCV tumor-adjacent samples, no differences were observed between HBV and HCV tumor samples.

**Figure 4:**
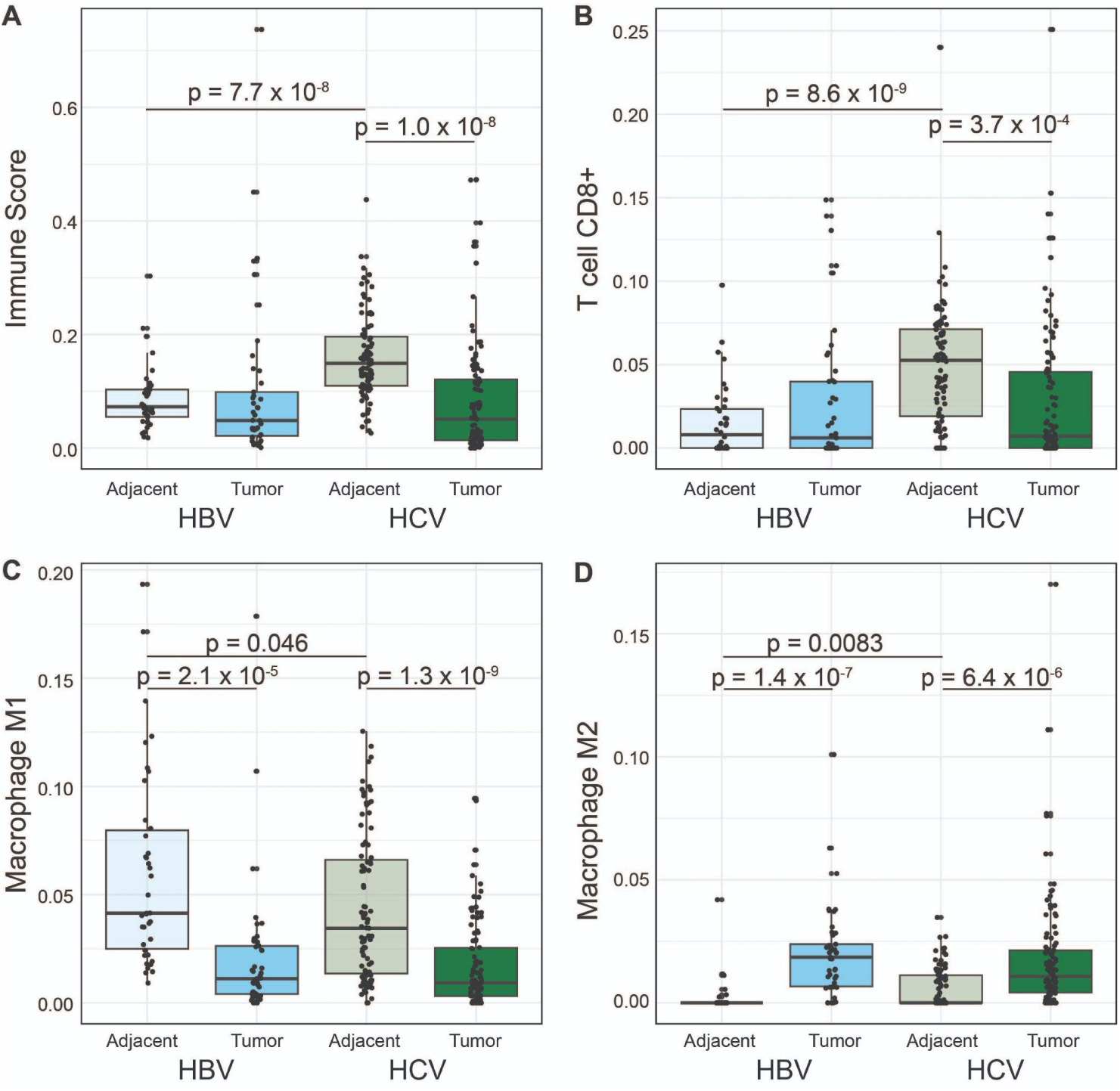
Greater immune infiltration in HCV tumor-adjacent tissue. **(A-B)** xCell immune deconvolution. **(A)** Comparison of the overall immune infiltration score in tumor and tumor-adjacent tissue segregated by etiology. **(B)** Comparison of CD8+ T cells in tumor and tumor-adjacent tissue segregated by etiology. **(D-E)** quanTIseq immune deconvolution, **(D)** comparison of M1 macrophages in tumor and tumor-adjacent tissue segregated by etiology, **(E)** comparison of M2 Macrophages in tumor and tumor-adjacent tissue segregated by etiology. For all boxplots, the bold line indicates the median and the upper and lower limits of the boxes indicate the 75th and 25th percentiles, respectively. The lower and upper whiskers indicate the minimum and maximum. Dots outside of the box and whiskers indicate outliers Significance tested with two-sample, two-sided t-tests.

Since immune infiltration is known to impact response to immunotherapy (30), we were interested in quantifying the relationship between immune cell infiltration in matched tumor-adjacent and tumor samples. We therefore classified each sample as either being above or below the median in terms of immune infiltration. We then classified paired samples into one of four groups: 1) groups with high tumor-adjacent immune infiltration that stayed high in tumor tissue, 2) groups with high tumor-adjacent immune infiltration that dropped to low infiltration in tumor tissue, 3) groups with low tumor-adjacent immune infiltration that stayed low in the tumor tissue, and 4) groups with low tumor-adjacent immune infiltration that changed to high in the tumor tissue (Supplementary Figure 3). Consistent with the high immune infiltration in HCV tumor-adjacent samples, the majority of HCV samples fell into the first two groups, with 29.5% (26/88) of samples having immune infiltration above the median in both tumor-adjacent and tumor samples and 52.3% (46/88) having immune infiltration above the median in only the tumor-adjacent samples. By contrast, HBV had only 10.5% (4/38) of samples having immune infiltration above the median in both tumor-adjacent and tumor samples and 34.2% (13/38) of samples having immune infiltration above the median in only the tumor-adjacent samples. Both HBV and HCV samples showed very low rates of samples where only the tumor sample had immune infiltration above the median (4.5%, 4/88 for HCV and 7.9%, 3/38 for HBV). The majority of HBV samples had low immune infiltration in both the tumor and tumor-adjacent samples (47.4%, 18/38) compared to a lower proportion for HCV samples (13.6%, 12/88).

We also performed immune cell deconvolution with quanTIseq (31) as a second approach. The results from quanTIseq confirmed increased CD8+ T cell infiltration in the HCV tumor-adjacent samples compared to all other samples (Supplementary Figure 2B). Additionally, deconvolution with quanTIseq demonstrated an increase in M2 polarized macrophages (Figure 4C, Supplementary Figure 2B) and a corresponding decrease in M1 polarized macrophages (Figure 4D, Supplementary Figure 2B) in tumor samples compared to non-tumor samples. Notably, the predominant M2 polarization was consistent across both HBV- and HCV-mediated liver cancer. The difference in macrophage polarization was not detected in the results from xCell, which may be attributable to the overlap in the M1 and M2 gene signatures for xCell. In the development of quanTIseq, marker genes were selected as a subset of the gene sets proposed for xCell with the removal of any signature genes included in the marker genes for a different cell type. Therefore, quanTIseq may be better equipped to identify differences in subpopulations of cells from a single progenitor cell that may share large numbers of marker genes. Overall, immune cell deconvolution demonstrated a strong immune cell infiltrate in the HCV tumor-adjacent tissue and a consistent immune profile between HBV and HCV tumor samples.

### Sex differential expression analysis highlights the need for increased sample size for female HCC

We next explored whether we could identify patterns of sex-differentially expressed genes within HBV and HCV-mediated liver cancer. We performed tumor:tumor-adjacent differential expression analyses, subset by sex and etiology (Figure 5). In HBV, we noticed a striking excess in the number of differentially expressed genes identified in the tumor:tumor-adjacent comparison in male patients that were not identified in the female subset (Figure 5A-C). We, therefore, repeated this analysis with random down-sampling of the male patients and demonstrated that the number of unique differentially expressed genes in males and females was approximately equivalent (Figure 5D). We repeated this approach to study sex differences in HCV-mediated liver cancer, and similarly observed a higher number of differentially expressed genes in the male tumor:tumor-adjacent analysis when sample sizes were unequal (Figure 5E-G) but not when we down-sampled to have equal representation among samples from males and females (Figure 5H).

**Figure 5:**
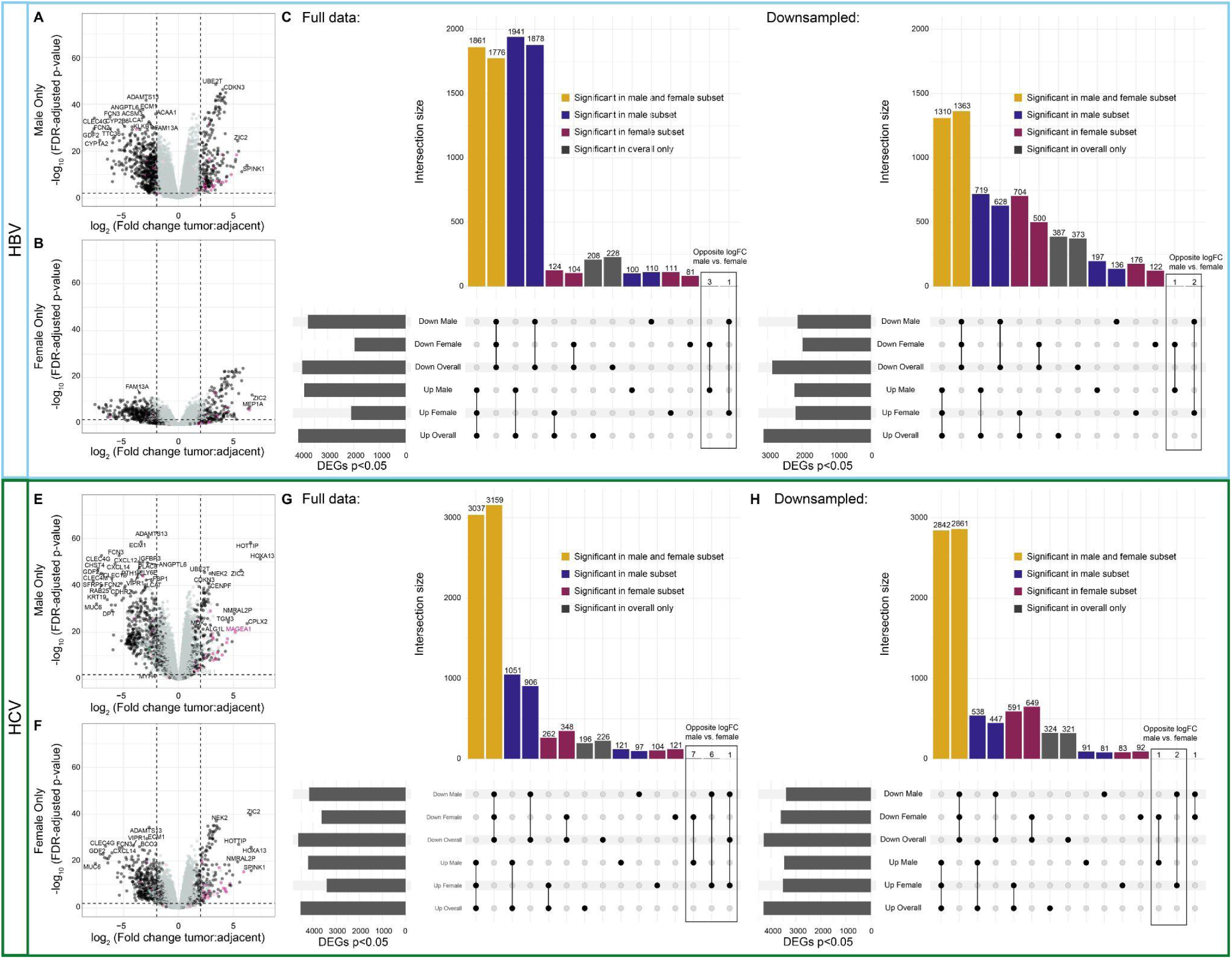
Tumor:tumor-adjacent differential expression in HBV and HCV samples is dominated by male differentially expressed genes. **A-B)** Volcano plots of differentially expressed genes from **(A)** male HBV samples and **(B)** female HBV samples. X-linked genes are indicated in pink, Y-linked in green, and autosomal in black. **(C)** Upset plot of differentially expressed genes from each of the comparisons with the full sample size. **(D)** Upset plot of differentially expressed genes from each of the comparisons with down-sampling of male samples to be equal to the number of female samples. Genes shared in all comparisons are colored yellow, genes unique to males are blue, and genes unique to females are maroon. **E-F)** Volcano plots of differentially expressed genes from **(E)** male HCV samples and **(F)** female HCV samples. **(G)** Upset plot of differentially expressed genes from each of the comparisons with the full sample size. **H)** Upset plot of differentially expressed genes from each of the comparisons with down-sampling of male samples to be equal to the number of female samples.

We then evaluated whether the genes identified in the segregated analyses showed similar expression changes from tumor-adjacent to tumor tissue in male and female samples, despite not passing multiple testing corrections in both sexes (Figure 6A-H). Within HBV, we fit a linear model on all genes shared in the tumor:tumor-adjacent comparison from males and females and found that our model had an R-squared value of 0.914, suggesting that 91.4% of the variability in the data is explained through the model (Figure 6A). Of note, within the genes unique to the male tumor:tumor-adjacent comparison, the R-squared value is still 0.667, suggesting that there is still a high degree of similarity between the overall logFCs in males and females (Figure 6B). Within the genes significant with down-sampling, the R-squared value remains high at 0.755. Within the genes unique to the female tumor:tumor-adjacent comparison, the R-squared value drops to 0.446 and is 0.432 among the genes significant with down-sampling (Figure 6C). Of the genes differentially expressed in opposite directions in male tumor:tumor-adjacent vs. female tumor:tumor-adjacent, 4/4 had an absolute logFC of less than 1.5, with 3/4 having an absolute logFC less than 1, suggesting that the biological impact of these differences may be low (Figure 6D).

**Figure 6:**
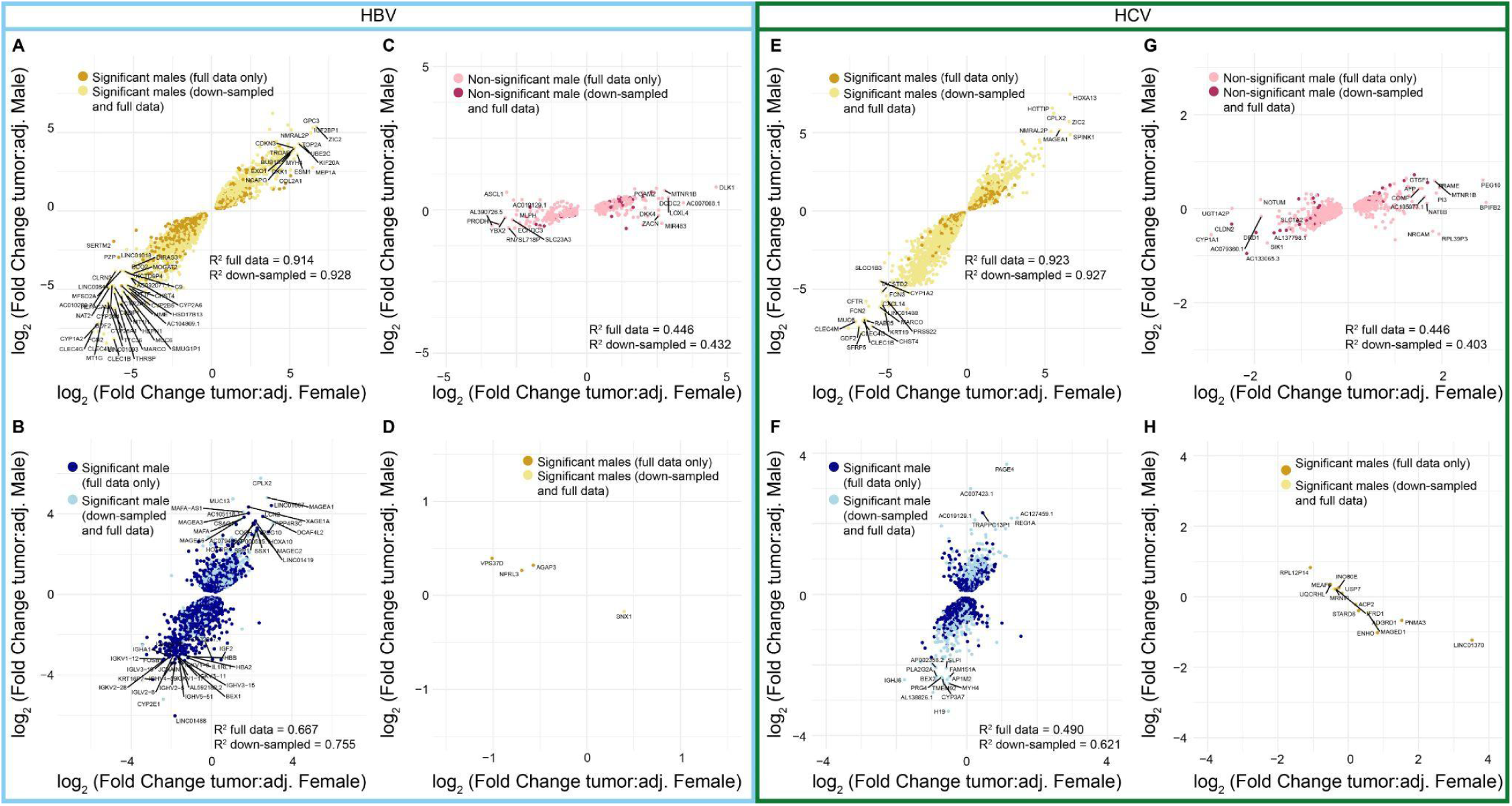
Comparisons of the log fold change values between males and females demonstrate consistency in direction and magnitude. Comparison of the logFC in males and females for **(A)** genes significant in both male and female tumors in the same direction in HBV, **(B)** genes significant only in male patients in HBV, **(C)** genes significant only in female patients in HBV, **(D)** genes significant in both male and female tumors in the opposite directions in HBV, **(E)** genes significant in both male and female tumors in the same direction in HCV, **(F)** genes significant only male patients in HCV, **(G)** genes significant only in female patients in HCV, and **(H)** genes significant in both male and female tumors in the opposite directions in HCV. For all plots, the lighter color genes indicate genes that were no longer significant (or gained significance for C/G) in males in the down-sampled differential expression analysis.

Within HCV, the genes shared in the tumor:tumor-adjacent comparison from males and females had an R-squared value of 0.923 (Figure 6E). Consistent with the larger sample sizes available for HCV, the genes unique to the tumor:tumor-adjacent comparison in males had an R-squared value of 0.490 that raised to 0.621 with down-sampling (Figure 6F). The genes unique to the tumor:tumor-adjacent comparison in females had an R-squared value of 0.447 that lowered slightly to 0.403 with down-sampling (Figure 6G). Finally, among the genes differentially expressed in opposite directions for the male tumor:tumor-adjacent and female tumor:tumor-adjacent comparisons, the majority still had logFCs less than 1.5, with only *LINC01370* and *PNMA3* demonstrating logFCs greater than 1.5 in females and none in males.

Finally, we performed a direct differential expression analysis between male tumor-adjacent and female tumor-adjacent tissue as well as between male tumor and female tumor tissue, subset by etiology (Supplementary Figure 4). Across all comparisons, the majority of the differentially expressed genes identified were on sex chromosomes, consistent with the expected sex differences. Of note, a larger number of autosomal genes were identified as differentially expressed between male HCV tumors and female HCV tumors, suggesting that there may be an increase in sex differences in these samples (Supplementary Figure 4D). Pathway enrichment analysis did not identify significantly enriched pathways in the male HBV tumor-adjacent:female HBV tumor-adjacent, male HBV tumor:female HBV tumor, or HCV:tumor-adjacent:female HCV tumor-adjacent comparisons. Within the HCV tumor:HCV tumor comparisons, several pathways were identified as enriched, including “Small molecule catabolic process”, “Diterpenoid metabolic process”, “Xenobiotic metabolic process” and “Hormone metabolic process” (Supplementary Figure 5).

## Discussion

Understanding the transcriptional and immune profile differences in HCC segregated by etiology and sex is critical to drive individualized therapeutic decision-making and rational design of therapeutic interventions in HCC. Towards this end, we performed differential expression and pathway analyses on HBV- and HCV-mediated HCC to probe the underlying etiology and sex differences.

Within the tumor:tumor-adjacent differential expression analysis segregated by viral etiology, we first demonstrated large changes in the genes identified in the overall tumor:tumor-adjacent comparison versus the tumor:tumor-adjacent comparisons subset by etiology. In particular, we noted the effect of sample size on the differential expression analyses. Our sample size for HCV was over double that of HBV, and this was reflected in a much higher consensus between the differential expression results for the overall tumor:tumor-adjacent comparison and the HCV-specific results compared to the HBV-specific results. This manifested as both a smaller number of genes shared between the overall and HBV-specific differential expression results and a larger number of differentially expressed genes identified only in the HBV-specific subset. These results highlight the importance of considering the composition of the samples when identifying differentially expressed genes in a mixed-etiology group.

Of particular interest, a small number of genes demonstrated significant log fold changes in opposite directions in the tumor:tumor-adjacent comparison for HBV compared to that for HCV. While the majority of these genes were identified at low fold changes, suggesting limited biological effects, a small number demonstrated large opposite fold changes. Among these candidates, several have previous associations with HCC pathogenesis in the literature, including *BEX2* (24,25), *AP1M2* (26), and *KRT23* (27,28). *BEX2* was identified as upregulated in HBV tumor tissue (logFC = 1.3, p = 0.0026) and down-regulated in HCV tumor tissue (logFC = -1.9, p = 7.6×10^-11^). High *BEX2* expression has been previously implicated in the maintenance of cancer stem cells and poor prognosis in HCC (24). Additionally, in mouse models of HCC, *BEX2* has been implicated in the oncogenic pathways mediating cell proliferation and metastasis (25). *AP1M2* was identified as upregulated in HBV tumor tissue (logFC = 0.95, p = 0.048) and downregulated in HCV tumor tissue (logFC = -1.92, p = 3.24187 cl:152810^-8^). Upregulation in HBV tumor tissue is consistent with previous studies showing that AP1M2 upregulation by HBV is implicated in the proliferation of liver cancer cells (26). *KRT23* was upregulated in HBV tumor tissue (logFC = 1.74, p = 0.022) and downregulated in HCV tumor tissue (logFC = -1.54, p = 0.00095). Previous work has suggested that knockdown of *KRT23* reduced HCC cell line proliferation and metastasis (27). Overall, this demonstrates that there are genes with possible biological implications that are differentially regulated in HBV and HCV tumors.

Pathway enrichment analysis on the differentially expressed genes in the HBV tumor:tumor-adjacent and HCV tumor:tumor-adjacent comparisons demonstrate that the majority of genes uniquely differentially expressed in HCV tumor:tumor-adjacent are enriched in immune pathways. By contrast, genes uniquely differentially expressed in the HBV tumor:tumor-adjacent genes are enriched in apoptotic pathways, consistent with literature suggesting that HBV plays a significant role in modifying apoptotic pathways in HCC (32). When we further probed the differences in HBV tumor:HCV tumor and HBV tumor-adjacent:HCV tumor-adjacent comparisons, we found that the tumor-adjacent samples had more differentially expressed genes than the tumor comparison. Pathway enrichment analysis on the tumor-adjacent comparison showed significant enrichment across multiple immune pathways.

Due to the repeated observation of enriched immune pathways, we probed the immune infiltration using immune deconvolution analyses. We demonstrated that the HCV tumor-adjacent samples had the highest immune infiltration, with higher immune infiltration than any of the other groups. Of note, despite the high immune infiltration in HCV tumor-adjacent compared to HCV tumor samples, no difference was observed in the immune infiltration in the HCV tumor compared to HBV tumor samples. This is consistent with prior work (17) and suggests that the tumor microenvironment is converging despite differences in the surrounding regions. We also demonstrated that HBV and HCV tumors had more M2 macrophages and fewer M1 macrophages than tumor-adjacent tissue, suggesting a more immunosuppressive and pro-tumorigenic immune environment, consistent with previous work (33,34). Of note, despite known sex differences in the immune function (21–23), there were no clear differences in the immune infiltration in tumor-adjacent or tumor samples between males and females in this dataset.

To probe how the immune infiltration in the tumor-adjacent tissue impacts the immune infiltration in the tumor tissue, we compared immune infiltration in paired samples. We demonstrated that over half of the HCV tumors have high immune infiltration in tumor-adjacent tissue that drops down in the tumor tissue. By contrast, the majority of HBV tumors have low immune infiltration in both the tumor-adjacent and tumor tissue. Immune infiltration has previously been identified as a marker of response to immunotherapy (30,35). However, this work raises an interesting question of how the immune infiltration in the tumor-adjacent tissue influences immune infiltration in the emerging tumor and how this impacts treatment efficacy, especially in the context of immunotherapy.

Finally, differential expression analyses in tumor:tumor-adjacent tissue segregated by sex were substantially limited by the low availability of female samples. Specifically, in HBV, the initial segregated analysis suggested a large predominance of genes significant in only males, but down-sampling of the male samples demonstrated that this observation was a feature of sample size rather than biology. Additionally, comparisons of the logFCs in males and females suggested that there was still a strong degree of association between the logFCs of genes that were only identified as significant in males, again reaffirming that the differences are likely attributable to sample size. While the differences in sample size between males and females prohibited extensive analysis of underlying biological differences in HCC between males and females, this analysis demonstrates the importance of increasing the available sequencing data in female HCC to enable a more extensive analysis of sex-specific differentially expressed genes.

Overall, we have demonstrated the importance of considering sex and viral etiology in future studies of HCC. HBV- and HCV-mediated HCC appear to converge on similar transcriptomic and immune profiles relative to the surrounding tissue. However, the impact of the differences in the tumor-adjacent tissue on treatment efficacy, particularly of immunotherapies, warrants further research. Additionally, understanding the profiles of the adjacent tissues may further illuminate the pathogenesis and immune evasion pathways in viral-mediated liver cancer. Preliminary data offered here demonstrate that there are sex differences in viral-mediated HCC that cannot be fully characterized with the limited sample sizes available. This finding highlights the importance of increased sampling of female tumors. Finally, the work on both viral- and sex-based differences in HCC underscores that the interpretation of differential expression analyses should be cautious in cases of unequal representation across groups.

## Methods

### Data acquisition and processing

Whole transcriptome data (RNA-seq) from tumor and tumor-adjacent viral-mediated HCC samples were obtained from the International Cancer Genome Consortium LIRI-JP dataset (controlled access permission to Dr. Kenneth Buetow via project DACO-1938) (36). Tumor-adjacent samples were taken from adjacent liver tissue. Healthy liver samples from unaffected individuals were analyzed from data collected from the GTEx consortium (controlled access permission to Dr. Melissa Wilson via project #36761: "Assessing shared and divergent sex differences across disease and healthy tissues"). All data analysis was performed with the Arizona State University High-Performance Computing resources (37). RNA sequencing FASTA files were visualized for quality using FASTQC (38). Samples from one individual (RK023) were excluded due to poor quality. Data was trimmed using Trimmomatic with parameters of 2 seed mismatches, palindrome clip threshold 30, simple clip threshold 10, leading quality value 3, trailing quality value 3, sliding window size 4, minimum window quality 30, and minimum read length of 50 (39). Transcript expression levels were quantified using Salmon (40). The Salmon index was built based on the Gencode HG38 version 29 genome. Pseudoalignment was carried out using automatic library-type detection. 147 (53.3%) of the 276 samples were detected by Salmon to be stranded, and 129 (46.7%) were unstranded. To verify this, we additionally used GuessMyLT (41) to infer the library type based on the FASTQ files. The results of this analysis are concordant with the library types detected by Salmon.

### Sample quality control

We utilized multidimensional scaling as implemented in the plotMDS function of the R package *limma* v. 3.40.2 to calculate distances between samples based on gene expression levels (42). The effect of the sequencing library type (stranded/non-stranded, Supplementary Figure 5A). After removal of the sequencing library type, the majority of samples clustered strongly by tumor/tumor-adjacent on principal component 1 and sex on principal component 2 (Supplementary Figure 5B). However, a small number of samples were found in clusters that did not match their annotation. For all samples, we plotted the gene expression of XIST, DDX3Y, UTY, and USP9Y (Supplementary Figure 5C-D), and six samples were excluded from subsequent analysis where the inferred sex chromosome complement from gene expression did not match the reported sex of the patient. Two paired samples had tumor and tumor-adjacent samples that clustered in the opposite groups (e.g., the sample marked as “tumor” grouped with “tumor-adjacent”, and the matched “tumor adjacent” sample grouped with “tumor” samples) and the annotations for these were changed to match the appropriate group. The distribution of the final cohort by tumor tumor-adjacent, sex, and etiology included 276 samples (Table 1).

**Table 1.** International Cancer Genome Consortium LIRI-JP dataset samples, segregated by sex and etiology. Samples included are those that passed all quality control steps.

|  | Reported Male Patient |  | Reported Female Patient |  |
| --- | --- | --- | --- | --- |
|  | Tumor | Tumor-Adjacent | Tumor | Tumor-Adjacent |
| HBV | 37 | 31 | 8 | 7 |
| HCV | 70 | 59 | 32 | 32 |
| **Total** | **107** | **90** | **40** | **39** |

### Filtering and Processing

FPKM (fragments per kilobase of exon per million fragments mapped) and TMM (Trimmed Mean of M-values) were obtained using *EdgeR* (43). We retained genes with a mean FPKM value of ≥ 0.5 in any group segregated by sex and viral status and read count of > 6 in at least 10 samples across all samples under investigation, resulting in 12,466 genes.

### Differential Expression

To detect differentially expressed genes between the sexes, tissues, and viral infection cases, we used linear regression as implemented in *limma*. Filtered, raw counts per million reads (CPM) were log2 normalized and adjusted for quality using the *voomWithQualityWeights* function in the limma R package (44). Differential expression analyses were performed for the following comparisons: (1) all tumor vs. all tumor-adjacent, (2) HBV tumor vs. HBV tumor-adjacent, (3) HCV tumor vs. HCV tumor-adjacent, (4) HBV tumor vs. HCV tumor, (5) HBV tumor-adjacent vs. HCV tumor-adjacent, (5) male HBV tumor vs. male HBV tumor-adjacent, (6) male HCV tumor vs. male HCV tumor-adjacent, (7) female HBV tumor vs. female HBV tumor-adjacent, (8) female HCV tumor vs. female HCV tumor-adjacent, (9) male tumor vs. female tumor, and (10) male tumor-adjacent vs. female tumor-adjacent. For the male tumor:tumor-adjacent and female tumor:tumor-adjacent comparisons, differential expression analysis was repeated with a randomly down-sampled set of samples to make the number of male and female samples equivalent. Library type was added as a covariate for each model. Correlation between measurements between tumor and tumor-adjacent samples from the same patient was accounted for in the linear modeling using the *duplicateCorrelation* function (42). Differentially expressed genes were identified using the *limma/voom* pipeline with empirical Bayes statistics. Upset plots were generated using the UpSetR package (45).

### Pathway Enrichment Analysis

Differentially expressed genes were then analyzed for overrepresentation of biological processes. Hypergeometric testing was performed using *clusterProfiler* (*46*) to identify significantly overrepresented gene ontology (GO) terms. P-values were adjusted for false discovery rate and GO terms with an adjusted p-value of less than 0.05 were considered as significantly overrepresented pathways. GO enrichment analyses were visualized as treemaps using ReviGo (47).

### Immune cell deconvolution

Immune cell deconvolution was performed across all samples with transcript per million (TPM) normalized expression data using xCell (29) and quanTIseq (31) through the *immunedeconv* R package (48).

### Data Access

Code for all analyses and visualizations performed is available here: https://github.com/SexChrLab/Viral_HCC_Sex_Diff.

## Author contributions

Conceptualization E.S.B., K.H.B., M.A.W. Methodology E.S.B., K.H.B., M.A.W. Formal Analysis E.S.B., A.J., H.M.N. Investigation E.S.B., A.J., H.M.N. Resources K.T.H, K.H.B, M.A.W. Writing -- Original Draft E.S.B., K.H.B., M.A.W. Writing -- Reviewing and Editing E.S.B., H.M.N., K.T.H., K.H.B., M.A.W. Visualization E.S.B., A.J. Supervision K.H.B. M.A.W.

## Acknowledgments

This publication was supported by the National Institute of General Medical Sciences of the National Institutes of Health under Award Number R35GM124827 to MAW. The content is solely the responsibility of the authors and does not necessarily represent the official views of the National Institutes of Health. ESB was supported by an F30 fellowship, F30CA281056. The authors acknowledge Research Computing at Arizona State University for providing high-performance computing resources that have contributed to the research results reported within this paper. We thank Dr. Tanya Phung for her assistance with data processing.

## Supporting Information Captions

**Supplementary Figure 1:**
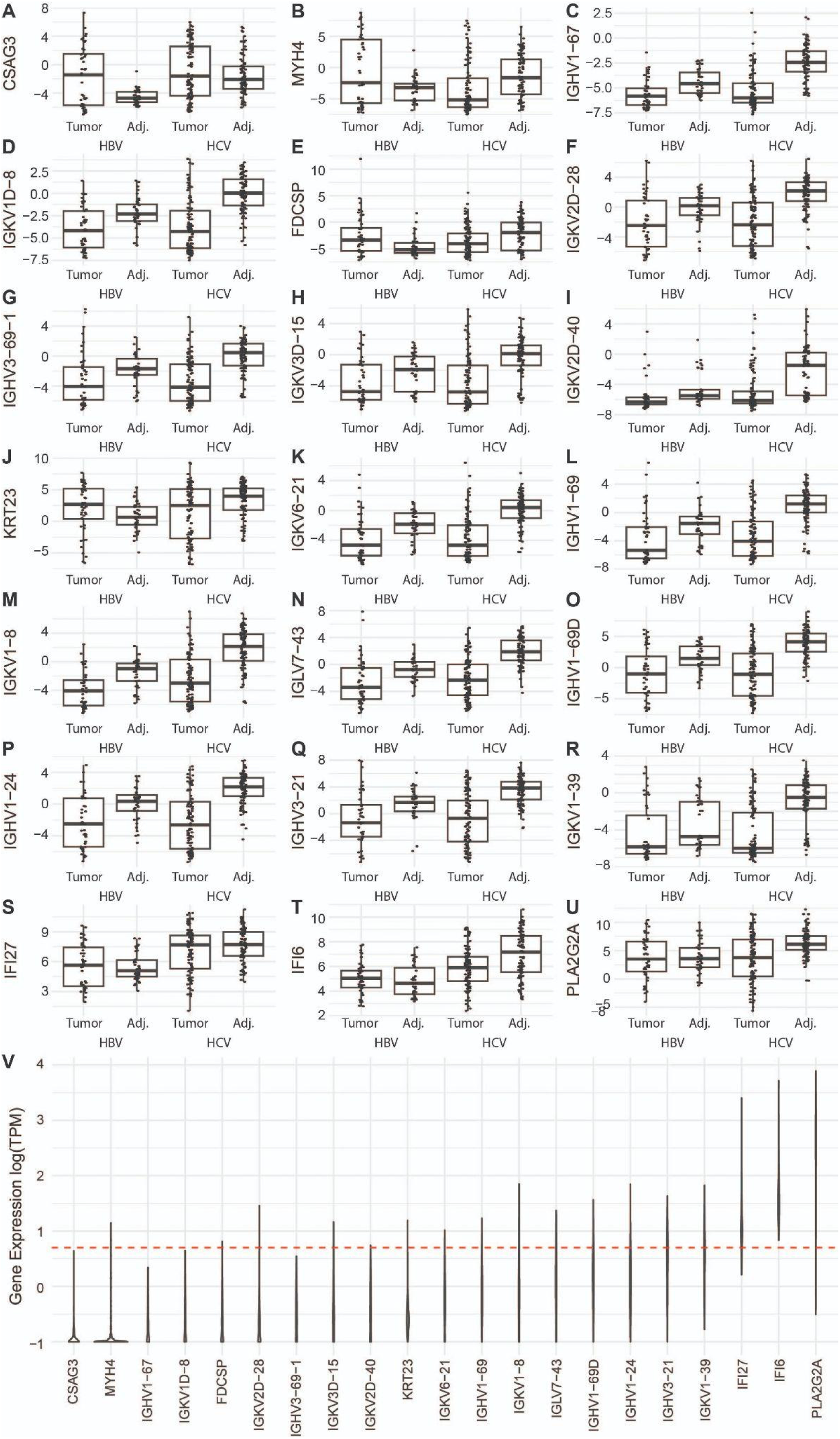
Expression of top differentially expressed genes from HBV tumor-adjacent to HCV tumor-adjacent tissue. **(A-U)** Boxplots demonstrating the *voom* normalized expression of each gene upregulated in HCV vs. HBV tumor-adjacent samples across tumor and tumor-adjacent, HBV and HCV samples. **(V)** Violin plot of the expression of genes upregulated in HCV tumor-adjacent tissue compared to HBV tumor-adjacent tissue in healthy liver tissue from the GTEx dataset. Expression in GTEx is normalized as transcripts per million (TPM) and is displayed on a log base 10 scale. The horizontal red dashed line corresponds to 1 TPM of gene expression.

**Supplemental Figure 2:**
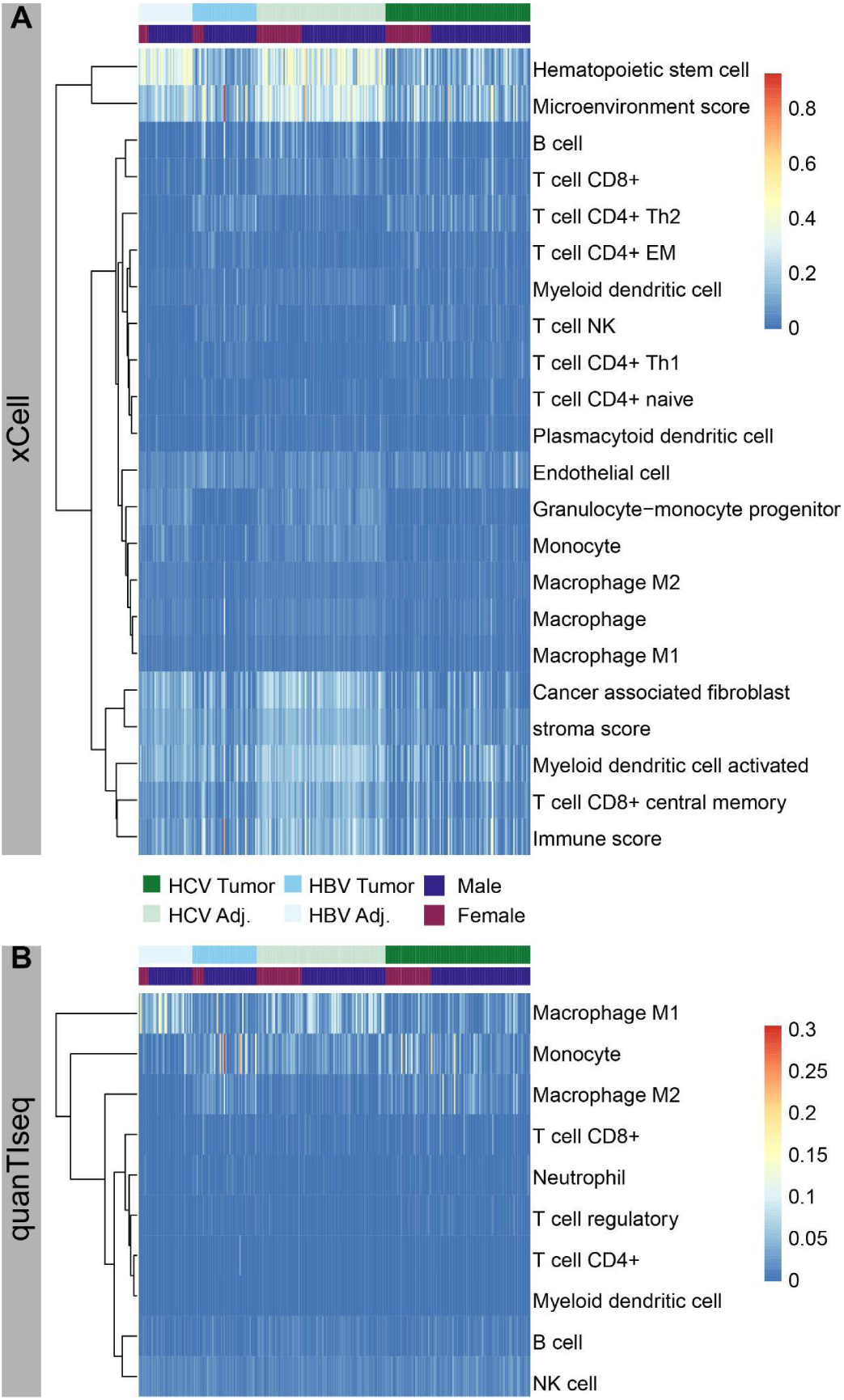
Full xCell and quanTIseq results across HBV and HCV samples. Heatmap of all immune cells identified with **(A)** xCell and **(B)** quanTIseq. Each column represents a single sample and the annotation bars across the top separate HCV tumor, HCV adjacent, HBV tumor, HBV adjacent, and male and female samples. Ajd.; adjacent.

**Supplemental Figure 3:**
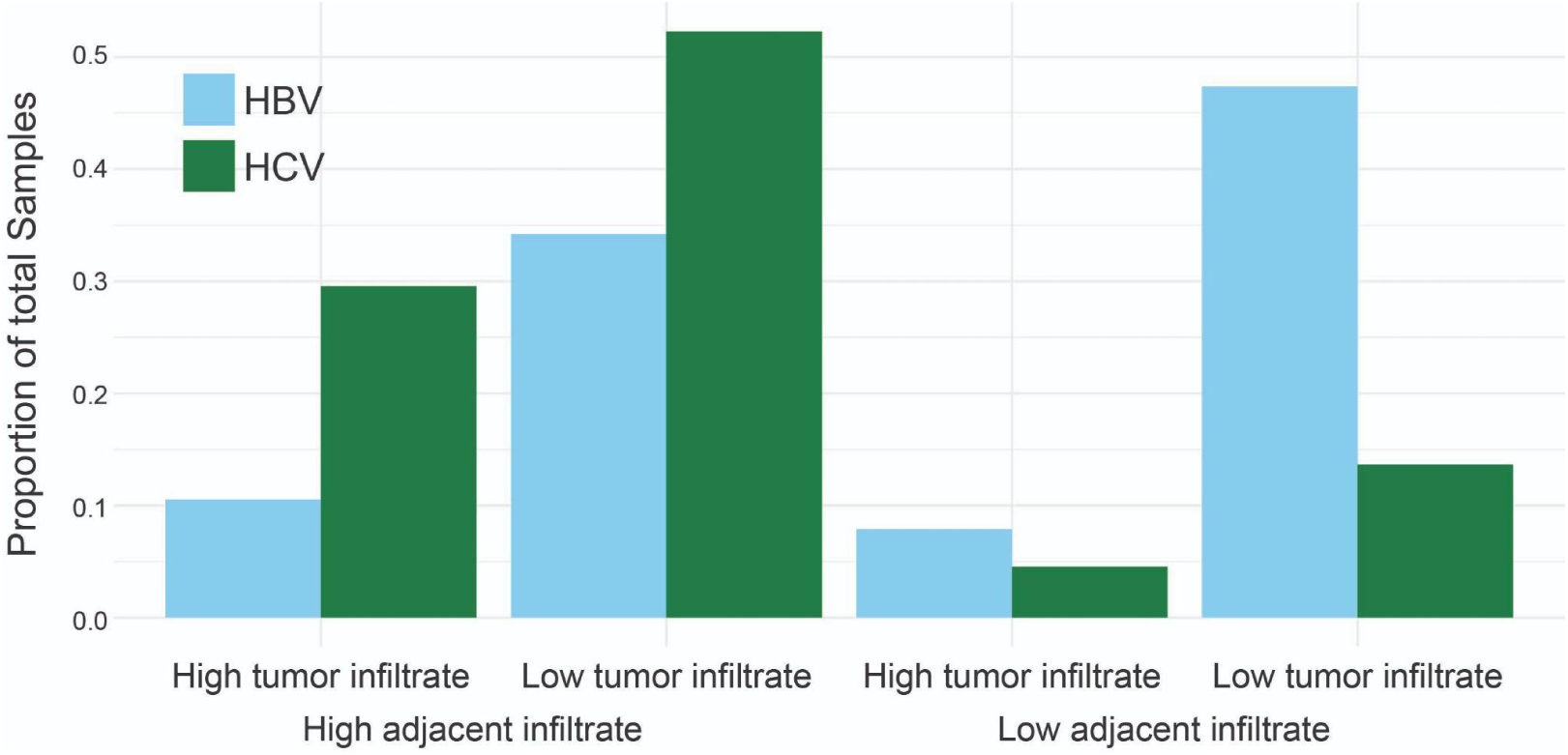
Comparison of infiltrate in tumor and adjacent matched samples. Comparison of the percent of HBV and HCV samples that fall into one of four categories of immune infiltration, high adjacent and tumor infiltration, high adjacent and low tumor infiltration, low adjacent and high tumor infiltration, and low adjacent and low tumor infiltration. High infiltration is defined as any infiltration above the median of all samples and low is defined as any infiltration below the median of all samples

**Supplementary Figure 4:**
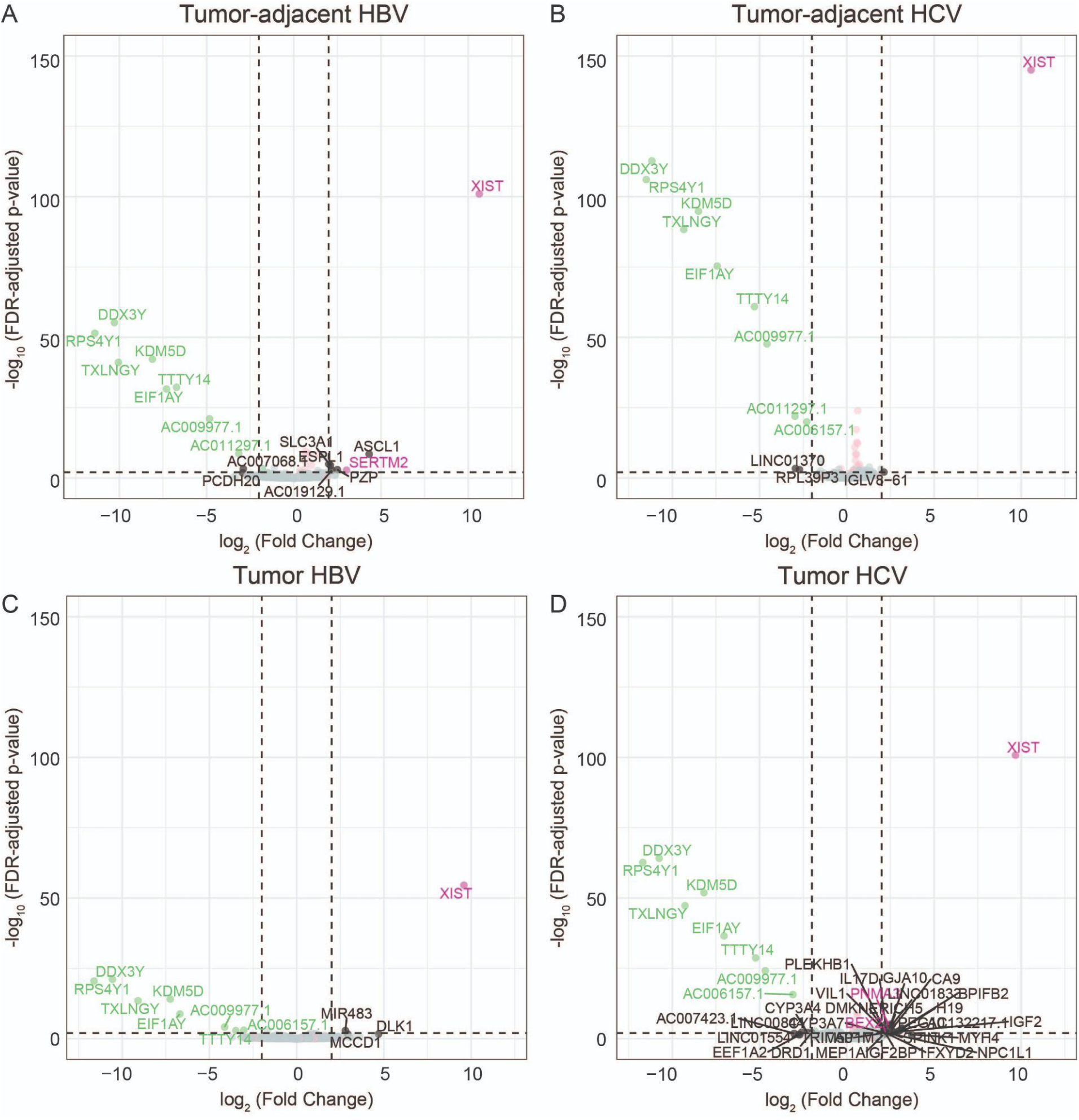
**A-D)** Volcano plots of differentially expressed genes from **(A)** male HBV tumor-adjacent:female HBV tumor-adjacent samples, **(B)** male HCV tumor-adjacent:female HCV tumor-adjacent samples, **(C)** male HBV tumor:female HBV tumor samples, and **(D)** male HCV tumor:female HCV tumor samples. X-linked genes are indicated in pink, Y-linked in green, and autosomal in black.

**Supplementary Figure 5:**
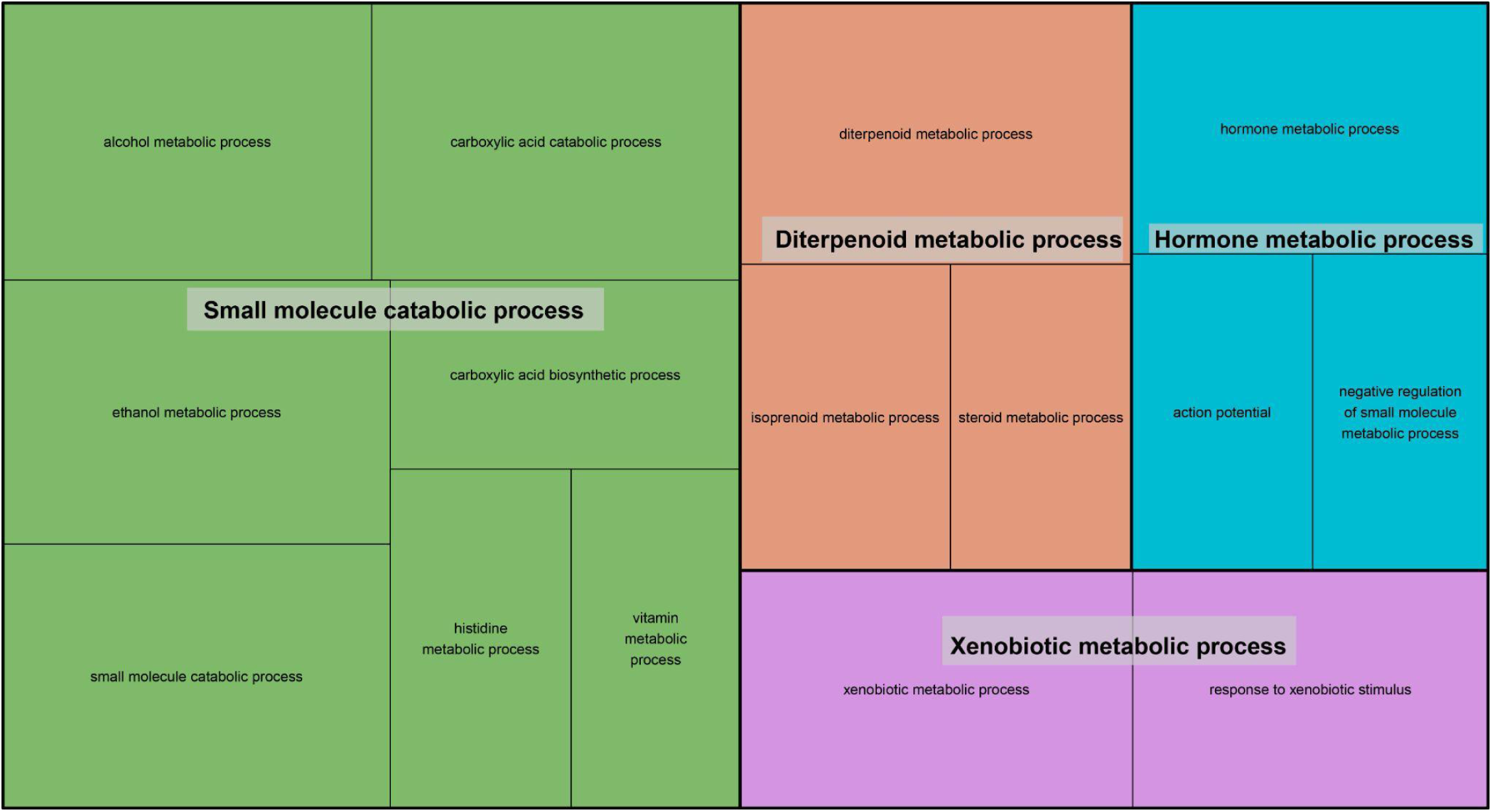
Pathways enriched in male HCV tumor:female HCV tumor comparison. Treemap visualization of GO enrichment analysis from all differentially expressed genes in the male HCV tumor:female HCV tumor comparison. The sizes of the boxes reflect the magnitude of the false-discovery adjusted p-value for the GO enrichment term.

**Supplementary Figure 5:**
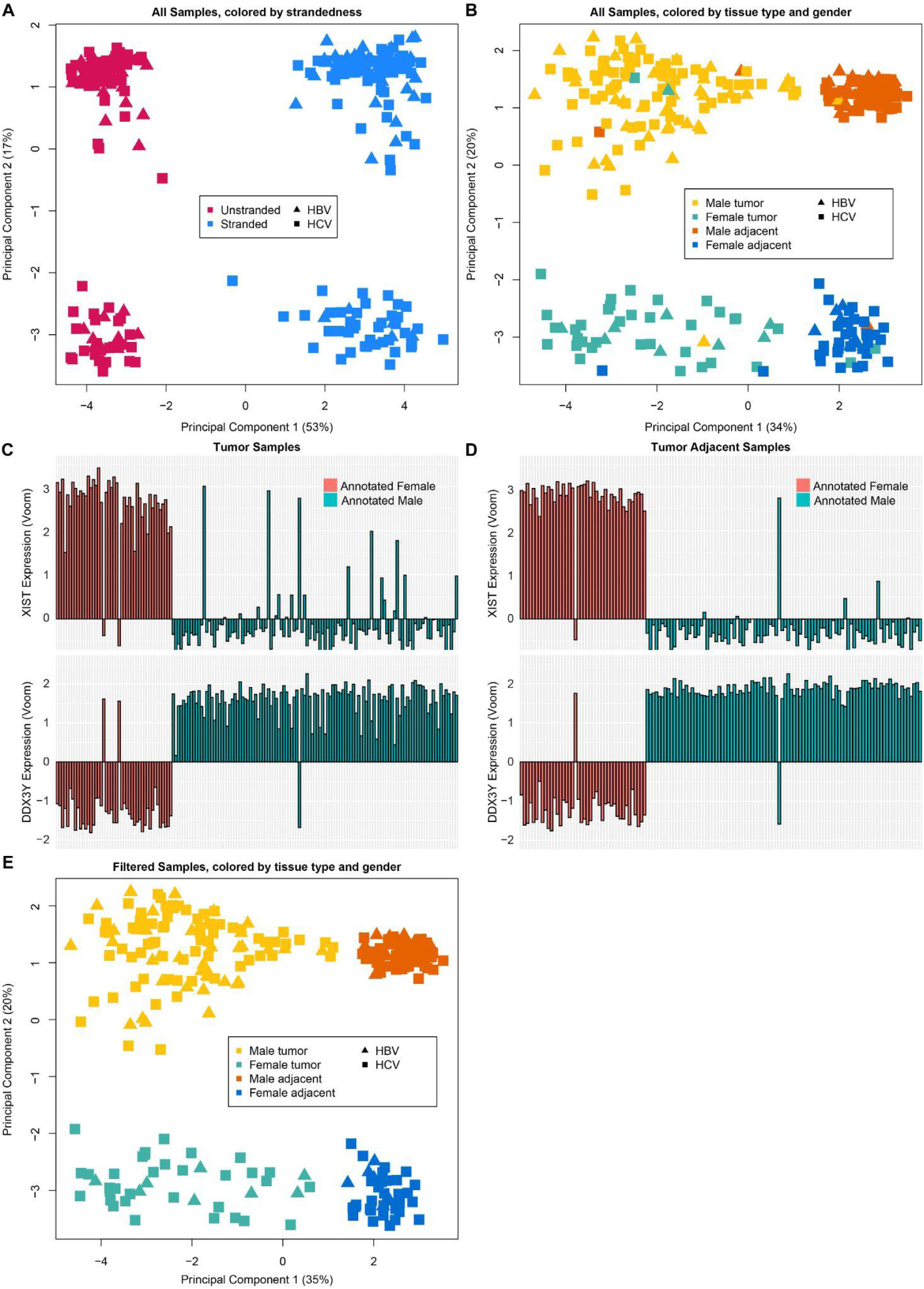
Quality control of all samples on tumor type and sex. MDS plot on the top 25 most variable genes, colored by **(A)** library type and **(B)** tumor status and sex. Plot of expression of XIST and DDX3Y across all **(C)** tumor samples and **(D)** tumor-adjacent samples. RK106, RK135, and RK105 were removed from subsequent analyses due to likely mislabeled sex supported by the MDS plots and expression of XIST and DDX3Y. RK066, RK113, and RK116 were removed due to the proximity of the paired sample suggesting that there may be cross-contamination of the samples. Finally, RK179 and RK065 had tumor and tumor-adjacent samples that were in opposite clusters on the MDS plot, therefore, these samples were relabeled to be consistent with their observed clusters. **(E)** MDS plot on the top 25 most variable genes colored by tumor status and sex.

